# Latent-based Directed Evolution accelerated by Gradient Ascent for Protein Sequence Design

**DOI:** 10.1101/2024.04.13.589381

**Authors:** Nhat Khang Ngo, Thanh V. T. Tran, Viet Thanh Duy Nguyen, Truong Son Hy

## Abstract

Directed evolution has been the most effective method for protein engineering that optimizes biological functionalities through a resource-intensive process of screening or selecting among a vast range of mutations. To mitigate this extensive procedure, recent advancements in machine learning-guided methodologies center around the establishment of a surrogate sequence-function model. In this paper, we propose Latent-based Directed Evolution (LDE), an evolutionary algorithm designed to prioritize the exploration of high-fitness mutants in the latent space. At its core, LDE is a regularized variational autoencoder (VAE), harnessing the capabilities of the state-of-the-art Protein Language Model (pLM), ESM-2, to construct a meaningful latent space of sequences. From this encoded representation, we present a novel approach for efficient traversal on the fitness landscape, employing a combination of gradient-based methods and directed evolution. Experimental evaluations conducted on eight protein sequence design tasks demonstrate the superior performance of our proposed LDE over previous baseline algorithms. Our implementation is publicly available at https://github.com/HySonLab/LatentDE.

## I. Introduction

THE field of protein engineering aims to strategically create or identify proteins that have practical applications with desired biological functions, such as fluorescence intensity [1], enzyme activity [2], and therapeutic efficacy [3]. The amino acid sequence of a protein governs the properties associated with its function through its spontaneous folding into three-dimensional structures [4], [5]. The mapping from protein sequence to functional property forms a protein fitness landscape [6] that characterizes the protein functional levels. The evolution in nature can be regarded as a searching procedure on the protein fitness landscape [7]. This natural process inspires the innovation of *directed evolution* [8]–[10], the most widely-applied approach for engineering protein sequences. It starts with a protein having some level of the desired function. Subsequently, a series of mutation and screening rounds are undertaken, wherein mutations are introduced to generate a collection of variant proteins. Through this iterative process, the most optimal variant is identified, and the cycle continues until a satisfactory level of improvement is achieved.

Recent machine learning (ML) methods have been studied to improve the sample efficiency of this evolutionary search [11]–[14]. However, these approaches require repetitive rounds of random mutagenesis and wet-lab validation, which are both money-consuming and time-intensive. In addition, while these methods can reduce experimental screening load by proposing potentially promising sequences, they only operate within a discrete space and make modifications to the amino acid sequence directly. In this manner, the challenge of navigating the sequence space remains unaddressed. The search space of protein sequences is discrete and combinatorially large, and most proteins have low fitness [15]. Moreover, protein fitness datasets are usually scarce as creating them requires costly wet-lab experiments [16]. These challenges make ML methods often stuck in local optima and prone to predict false positive samples [17]. An alternative methodology to working in the sequence space is to focus on acquiring a low-dimensional, semantically rich representation of proteins. These latent representations collectively form a latent space, offering more navigable and smoother terrain as they encapsulate the fitness landscape of the sequences into continuous forms. With this approach, a therapeutic candidate can be optimized using its latent representation in a procedure called latent space optimization (LSO). However, the integration of LSO with directed evolution remains a relatively unexplored territory.

Designing high-fitness protein sequences is regarded as a black-box model-based optimization (MBO) problem, where the primary goal is to explore design inputs maximizing a black-box objective (e.g., fitness). Typically addressed through an online iterative process, online MBO, in each iteration, proposes novel designs and solicits feedback from the unknown objective function to enhance subsequent design proposals. However, accessing the true objective function of protein fitness proves challenging due to constraints such as limited experimental data and computational resources. On the other hand, offline MBO emerges as an efficient for design desirable outputs without accessing true objection functions. Offline algorithms are allowed to observe a static dataset of designs and produce data samples that are later re-evaluated by the truth objective function. Albeit efficient for optimization, offline MBO often faces an out-of-distribution issue wherein the optimized design input *x* may be unseen to a learned proxy 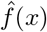, which replaces the ground truth objective for evaluating the design outputs. As 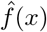 is trained to approximate the fitness landscape of a training dataset, out-of-distribution inputs may negatively affect its predictions [18], thereby fooling the discovery algorithm to produce non-desirable designs.

To overcome the above challenges, in this paper, we propose **L**atent-based **D**irected **E**volution (LDE), a novel approach that, to the best of our knowledge, is the first latent-based method for directed evolution. Specifically, by leveraging the latent space of ESM-2 [19], a state-of-the-art pre-trained protein language model (pLM), LDE learns to reconstruct and predict the fitness value of the input sequences in the form of a variational autoencoder (VAE) regularized by supervised signals. Post-training, LDE addresses the limitations of the two MBO methods by combining the best of both worlds. Starting from a wild-type sequence, LDE first encodes it into the latent representation, on which the gradient ascent is performed as an efficient offline MBO algorithm that guides the latent codes to reach high-fitness regions on the simulated landscape. Within these regions, to address the out-of-distribution limitation of the offline counterpart, LDE integrates latent-based directed evolution. This involves iterative rounds of randomly adding scaled noise to the latent representations, facilitating local exploration around high-fitness regions. The noised latent representations are decoded into sequences and evaluated by the truth oracles (e.g., wet-lab experiments, large ML models), and top samples are selected for the next round of the algorithm, enabling an efficient gradient-free sampling strategy. In summary, the contributions of our paper are outlined as follows:

- We propose a latent-based method for directed evolution for biological sequence design.
- We address the limitations of offline and online MBO methods by combining the best of both worlds, using gradient ascent as the warm-up procedure to guide the latent representations to feasible regions for accelerating latent-based directed evolution.
- Empirical evaluations conducted on eight protein datasets demonstrate that our proposed LDE surpasses the SOTA method in terms of fitness scores. We additionally conduct several ablation studies to showcase the effectiveness of our method, as well as to gain more insights into ML-based protein design.

## II. Related Works

### A. ML for Fitness Landscape Modeling

The application of ML in protein engineering has experienced a significant upswing, particularly in the domain of modeling the protein fitness landscape. Several methodologies have been proposed, leveraging co-evolutionary information extracted from multiple sequence alignments to predict fitness scores [20]–[22]. Alternatively, pre-trained language models have been employed for transfer learning or zero-shot inference [23]–[26]. The learned protein landscape models can be used to replace the expensive wet-lab validation to screen enormously designed sequences [27], [28].

### B. Latent-based Methods for Sequence Design

Directed evolution represents a classical paradigm in protein sequence design that has achieved significant successes [10], [29]–[31]. Within this framework, various ML algorithms have been proposed to improve the sample efficiency of the evolutionary search [11], [13], [14], [27], [32]. However, the majority of these methods are designed to optimize protein sequences directly in the sequence space, dealing with discrete, high-dimensional decision variables. Alternatively, [33] and [34] employed a VAE model and applied gradient ascent to optimize the latent representation, which is subsequently used to decode into string-based representations of small molecules. Similarly, [35] employed an off-policy reinforcement learning method to facilitate updates in the representation space. Another notable approach [36] involves training a denoising autoencoder with a discriminative multi-task Gaussian process head, enabling gradient-based optimization of multi-objective acquisition functions in the latent space of the autoencoder. Recently, latent diffusion has been introduced for designing novel proteins, harnessing the capabilities of a protein language model [37]. Diverging from the aforementioned works, we introduce a novel approach wherein, within the latent representation of a VAE model, gradient ascent is employed as a warm-up phase for latent-based directed evolution.

## III. Preliminaries

### A. Directed Evolution Theory

Directed evolution (DE) is a conventional approach in protein engineering that aims to search for global maximal protein sequences from a large database of unlabeled candidates *S* with minimal experimental validation [38], [39]. DE employs an accelerated cycle of mutation and selection, which iteratively generates a pool of protein variants and selects those having improved desired properties as the next generation. In essence, directed evolution is a local exploration around the regions of high-fitness proteins in large sequence space. The algorithm typically starts with wild-type protein sequences and repeatedly mutates a relatively small number of their amino acids to obtain variants with enhanced properties. This is motivated by the observation of [7] showing that functional protein sequences are clustered into groups in a vast space of non-functional ones, making locally exploring around the seed sequences possible to find ones with improved functions. Indeed, we illustrate this phenomenon in the left part of Figure 1.

**Fig. 1.**
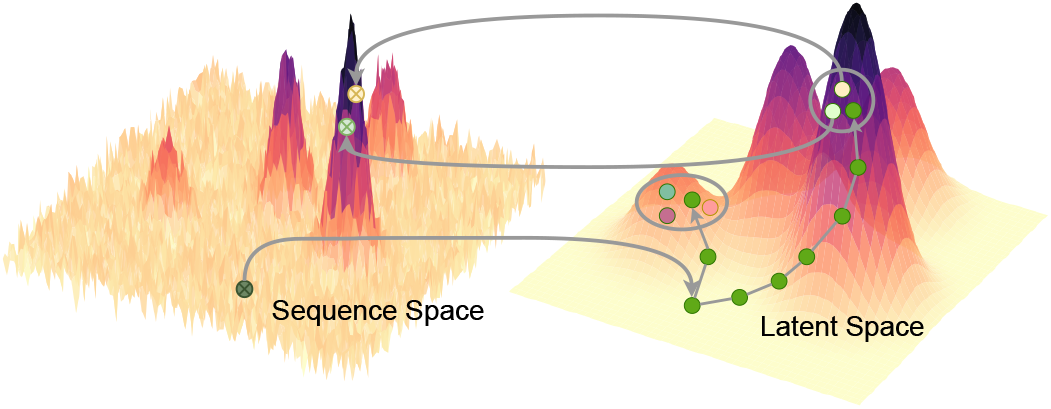
Optimizing protein fitness directly in sequence space is notoriously difficult due to its non-smooth and sparse landscape. We overcome this obstacle by performing directed evolution within the smooth and continuous latent space of a generative model. Our algorithm begins by identifying high-fitness regions through gradient ascent. Within these regions, we strategically sample a defined number of neighboring points (represented by gray circles) to create a diverse population for the evolutionary process. This population then undergoes iterative selection and mutation within the latent space, ultimately converging to sequences with enhanced fitness.

### B. Latent Space Optimization

Latent space optimization (LSO) is a model-based optimization technique that performs in the latent space *Ƶ* of deep generative models. In particular, an objective function *f* : *Ƶ*⟼ ℝ is trained to predict the objective values of data samples directly from their latent representations. When *Ƶ* is a low-dimensional and continuous space, LSO acts as an efficient and effective optimization method as it overcomes the difficulties of optimizing in discrete and high-dimensional spaces by turning the problem into the continuous version, which is simpler to solve with a wide range of available techniques like gradient ascent and Bayesian optimization (BO) [40]. Additionally, *f* can be trained by using an encoder *ϕ* : *𝒳* ⟼ *𝒱* that maps the input sample *x* ∈ *𝒳* to its corresponding latent point *z* ∈ *Ƶ*.

## IV Method

Optimizing the latent variables of generative models has proven effective in various drug and protein design tasks [36], [41]. Our work introduces a novel approach that utilizes directed evolution to efficiently discover optimal protein sequences directly from their latent representations.

We begin with the problem formulation in Section IV-A. Then, we present our pre-trained regularized variational autoencoders (VAEs) [42] in Section IV-B and how to perform directed evolution, which is accelerated by gradient ascent, in the latent space of VAEs in Section IV-C.

### A. Problem Formulation

We consider the problem of designing protein sequences to search for high-fitness sequences *s* in the sequence space *𝒱*^*L*^, where *𝒱* denotes the vocabulary of amino acids (i.e. |*𝒱*| ≈ 20 because both animal and plant proteins are made up of about 20 common amino acids) and *L* is the desired sequence length. We aim to design sequences that maximize a black-box protein fitness function *𝒪* : *𝒱*^*L*^ ⟼ ℝ which can only be measured by wet-lab experiments:

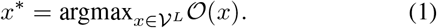

For in-silico evaluation, given a static dataset of all known sequences and fitness measurements *𝒟*^*\**^ = {(*x, y*)|*x* ∈ 𝒱^*L*^, *y* ∈ ℝ}, an oracle *𝒪*_*ψ*_ parameterized by *ψ* is trained to minimize the prediction error on *𝒟*^*\**^. Afterward, *𝒪*_*ψ*_ is used as an approximator for the black-box function *𝒪* to evaluate computational methods that are developed on a training subset *𝒟*_*t*_ of *𝒟*^*\**^. In other words, given *𝒟*_*t*_, our task is to generate sequences 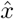 that optimize the fitness approximated by *𝒪*_*ψ*_.

In this paper, we focus on designing sequences upon wild-type sequences as the same with traditional evolution and exploration algorithms described in [9], [10], [27], [43]. Specifically, the algorithms commence with a single wild-type sequence and iteratively generate candidates with enhanced properties from previous references. Figure 2 illustrates the overview of our method.

**Fig. 2.**
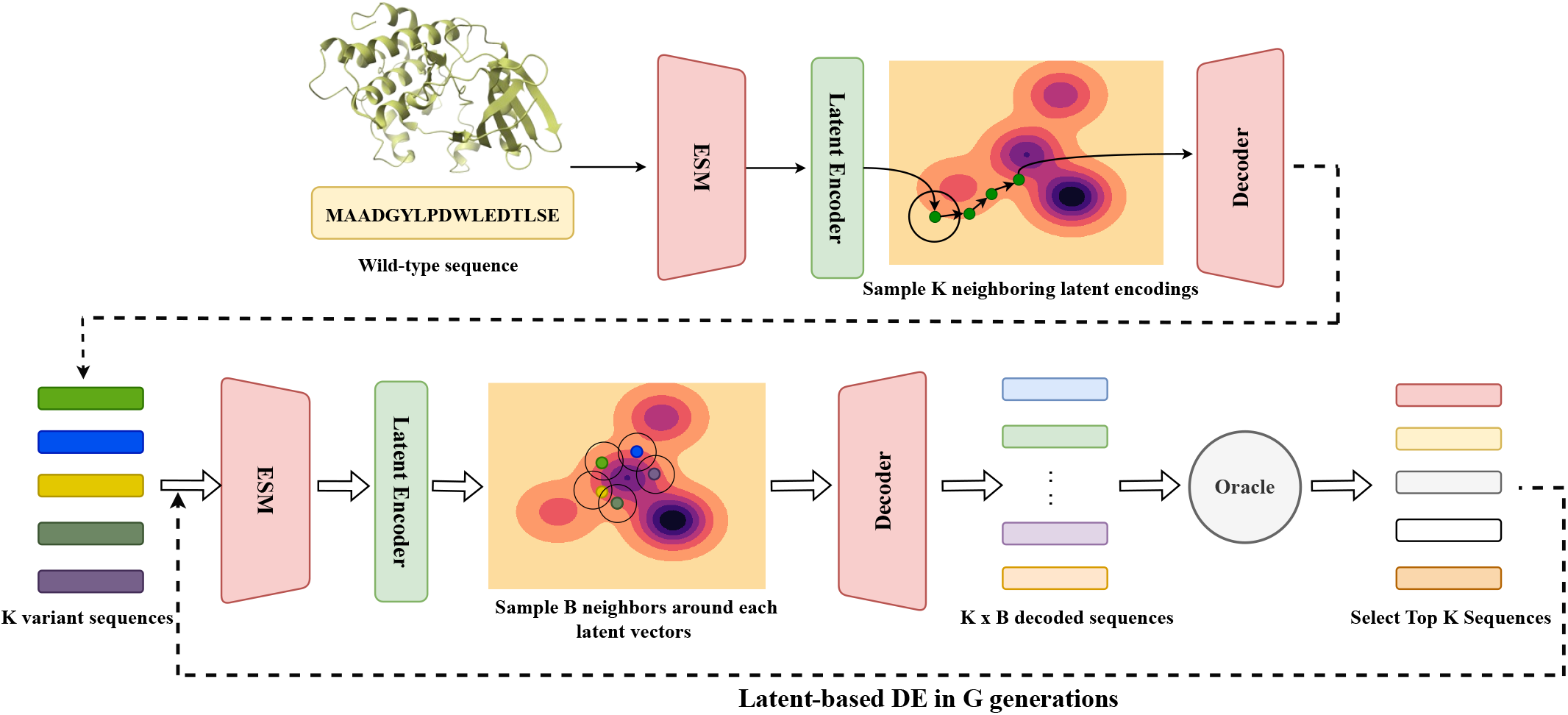
Overview of our proposed method. **Top**: We iteratively encode the wild-type sequence into latent variables and run gradient ascent to move them towards regions of high fitness on the approximated landscape. This process yields an initial population of K-diverse variant sequences. **Bottom** From *K* sequences, our latent-based directed evolution is performed in *G* generations. Sequences in the previous generation are encoded into the latent space at each generation. Then, *B* neighbors are sampled around those latent variables and decoded into *B* sequences, which are evaluated by a black-box oracle. The protein used in this figure is only for illustration purposes.

### B. Regularized Variational Autoencoder

Considering a dataset of protein sequences *x* ∈ 𝒱^*L*^ with corresponding fitness values *y* ∈ ℝ, denoted as 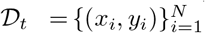, we train a VAE comprising an encoder 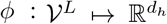 and a decoder *θ* : ℝ^*d*^ 1⟼ *Ƶ*^*L*^. Each sequence *x*_*i*_ is encoded into a low-dimensional vector 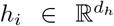. Subsequently, the mean *μ*_*i*_ ∈ ℝ^*d*^ and log-variance log *σ*_*i*_ ∈ ℝ^*d*^ of the variational posterior approximation are computed from *h*_*i*_ using two feed-forward networks, 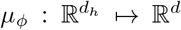 and 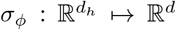. A latent vector *z*_*i*_ ∈ ℝ^*d*^ is then sampled from the Gaussian distribution 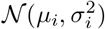, and the decoder *θ* maps *z*_*i*_ back to the reconstructed protein sequence 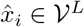. The training objective involves the cross-entropy loss 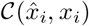 between the ground truth sequence *x*_*i*_ and the generated 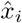, as well as the Kullback-Leibler (KL) divergence between 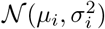 and *𝒩* (0, *I*_*d*_):

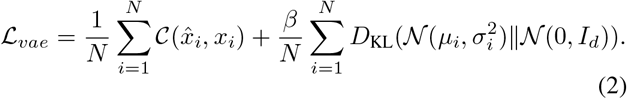

Here, *β* is the hyperparameter to control the disentanglement property of the VAE’s latent space.

#### a) Latent Space Regularization

One can consider the KL-divergence loss term in Equation (2) as a regularizer for training autoencoders. In particular, the KL-divergence regularizer makes the encoder *ϕ* produce latent space that is close to the unit Gaussian prior (i.e., the global optimum). The learned latent space could be smooth and convex, which is useful for model-based searching algorithms during optimization. However, forcing the approximate posterior *q*_*ϕ*_(*z*|*x*) to be unit Gaussian may lead to the fact that the encoder network *ϕ* may encode zero information into the latent space [44], resulting in less meaningful latent representations. This is not a good phenomenon for our latent-based directed evolution algorithm that aims at locally searching for high-fitness proteins by iteratively modifying the reference sequences (see Section III). Regarding learning protein fitness landscape, the percentage of high-fitness sequences in datasets is relatively small compared to those with low fitness scores. Consequently, the latent representations of high-fitness proteins may be hidden within a dense of low-fitness ones [34], leading to inefficient searching processes. To address this issue, following [34], we jointly train our VAEs with a fitness predictor *f* : ℝ^*d*^ ⟼ ℝ that maps the latent vectors *z ∈* ℝ^*d*^ to the fitness space. This leads to the final loss for training *ϕ, θ*, and *f* as:

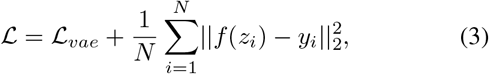

where *y*_*i*_ ∈ ℝ is the experimental fitness of sequence *x*_*i*_ *∈ 𝒱*^*L*^, which is encoded into latent vector *z*_*i*_ = *ϕ*(*x*_*i*_) ∈ ℝ^*d*^. This additional *L*_2_-regularizer forces the VAE’s encoder to generate latent representations of protein sequences with comparable fitness scores to be closely positioned in the latent space, thereby separating those with high fitness values from dense clouds associated with low-fitness regions.

### C. Latent-Based Directed Evolution

Inspired by nature’s evolutionary process, where subtle mutations in a protein’s sequence can significantly improve its fitness, our method explores the local region of the wild-type sequence’s latent encoding through iterative sampling from the approximate distribution encoded by the pretrained encoder *ϕ*. A crucial challenge arises from the possibility that the latent representation *z* of *x*^*wt*^ may be situated in low-fitness regions of the protein landscape. This can lead to slow convergence in directed evolution, where offspring generated by random mutations of low-fitness parents tend to inherit similar limitations. To address this problem, we incorporate gradient ascent as a preliminary step to generate the initial population *𝒫*_0_, accelerating the evolutionary process towards more advantageous regions. Algorithm 1 provides a detailed, line-by-line breakdown of our approach.

#### Algorithm 1

Latent-based directed evolution accelerated by gradient ascent

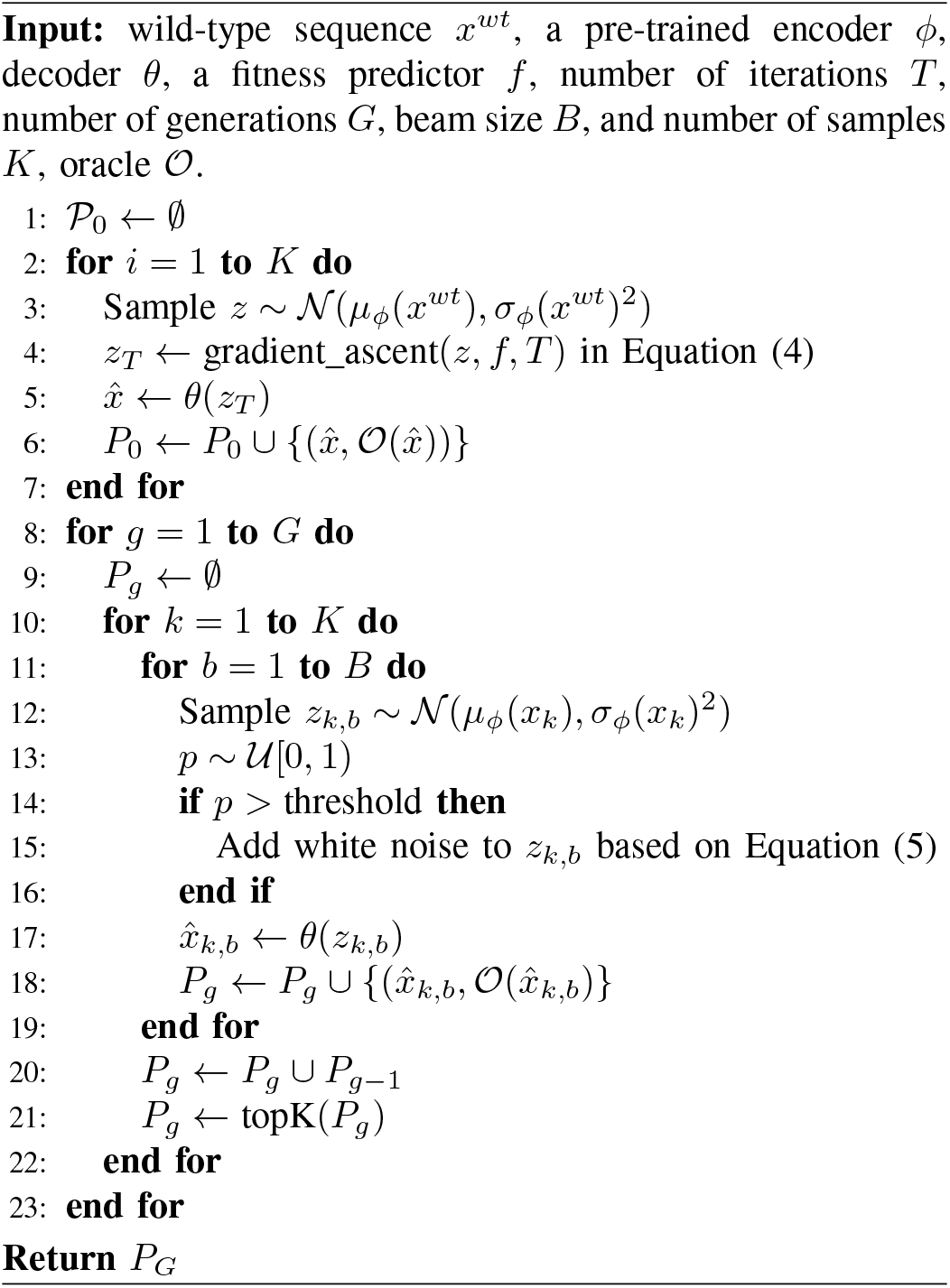

#### a) Gradient Ascent (GA)

At the beginning of our algorithm, the wild-type protein sequence *x*^*wt*^ is encoded by the encoder *ϕ* to produce its mean *μ*_*ϕ*_(*x*^*wt*^) and log variance log *σ*_*ϕ*_(*x*^*wt*^). A latent representation *z* is then sampled from *𝒩* (*μ*_*ϕ*_(*x*^*wt*^), *σ*_*ϕ*_(*x*^*wt*^)^2^). Subsequently, we perform gradient ascent with a learning rate of *α* in *T* iterations to move *z* to high-fitness regions as:

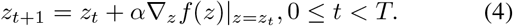

Where 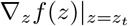 is the gradient of the fitness predictor *f* with respect to *z*_*t*_. After *T* iterations, we add *z*_*T*_ to the initial population *𝒫*_0_. The procedure is iteratively executed until *K* pairs of decoded sequences along with their respective fitness scores are obtained. At this stage, the fitness scores are computed by the latent predictor *f*, allowing fast computation and efficient latent sampling (see lines 1 to 7).

#### b) Evolutionary Process

From lines 8 to 23, a latent-based directed evolution is conducted in *G* generations to generate *K* protein sequences with high fitness values. For each candidate *x*_*k*_, we sample *B* latent variables from its posterior distribution, where each is denoted as *z*_*k,b*_ ∼ *𝒩* (*μ*_*ϕ*_(*x*_*k*_), *σ*_*ϕ*_(*x*_*k*_)^2^). These latent representations are then decoded into sequences by the decoder *θ*, i.e. 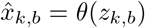. Wet-lab experiments then evaluate the decoded sequences to obtain their fitness scores. All sequences generated in generation *g* are added into *𝒫*_*g*_. Finally, at the end of *g*, the top *K* sequences are selected from both *𝒫*_*g*−1_ and *𝒫*_*g*_.

#### c) Random Exploration

Although sampling around the local areas of high-fitness latent codes can guarantee the superiority of the generated sequences, the search process may be prone to be trapped in these local regions after a certain number of generations, thereby hindering the exploration of potentially promising sequences, which are unseen before. As a result, we randomly add white noise to the latent variables when *p* ∼ *𝒰* [0, 1) is higher than a threshold. As demonstrated in line 14 of Algorithm 1, the formula is defined as:

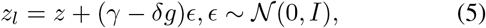

where *γ ∈* ℝ denotes the step size that controls the exploration rate, and *δ* is the annealing factor at the generation *g* of the evolution process. We hypothesize that when *g* gets close to *G*, i.e., the total number of generations, the population tends to contain superior samples; thus, we slow down the exploration to avoid degeneration at the end of the algorithm.

#### d) Efficient Sampling and Optimization

Directed evolution is an effective method in protein design. Yet, it requires intensive computational resources to directly mutate protein sequences that contain hundreds of amino acids. Our latent-based algorithm can improve the sampling efficiency in two perspectives. First, via VAEs, we can compress long protein sequences into continuous low-dimensional latent representations. This allows us to model the mutation process as a simple noise addition within the VAEs’ latent space. Compared with sophisticated mutation operators designed in discrete sequence spaces as in [9], [10], [32], [38], our approach is both simpler (requiring no domain knowledge) and more computationally efficient. The latent-based DE approach further unlocks the power of parallel computing on GPUs, a cornerstone of modern deep learning. This parallelization translates to dramatic efficiency gains, as mutation and sequence generation can be processed concurrently. Leveraging the compact nature of tensor-stored latent representations on GPUs dramatically accelerates these operations, making our algorithm well-suited for large-scale protein design tasks.

## V. Experiments

In this section, we undertake a comprehensive set of experiments to assess and validate the efficacy of our proposed LDE on the task of protein sequence design.

### A. Experimental Setup

#### a) Datasets

Following [27] and [28], we assess the performance of our method across eight protein engineering benchmarks: (1) Green Fluorescent Protein (**avGFP**), (2) Adeno-Associated Viruses (**AAV**), (3) TEM-1 *β*-Lactamase (**TEM**), (4) Ubiquitination Factor Ube4b (**E4B**), (5) Aliphatic Amide Hydrolase (**AMIE**), (6) Levoglucosan Kinase (**LGK**), (7) Poly(A)-binding Protein (**Pab1**), (8) SUMO E2 Conjugase (**UBE2I**). The comprehensive dataset information, including the protein name, organism, optimization target, sequence length, data size, and data percentiles, is provided in Table I. Each dataset represents distinct optimization target, which we simplify as a singular term *“fitness”* when reporting the benchmark results. Detailed descriptions of the data are provided in the Supplementary Material.

**TABLE I.**
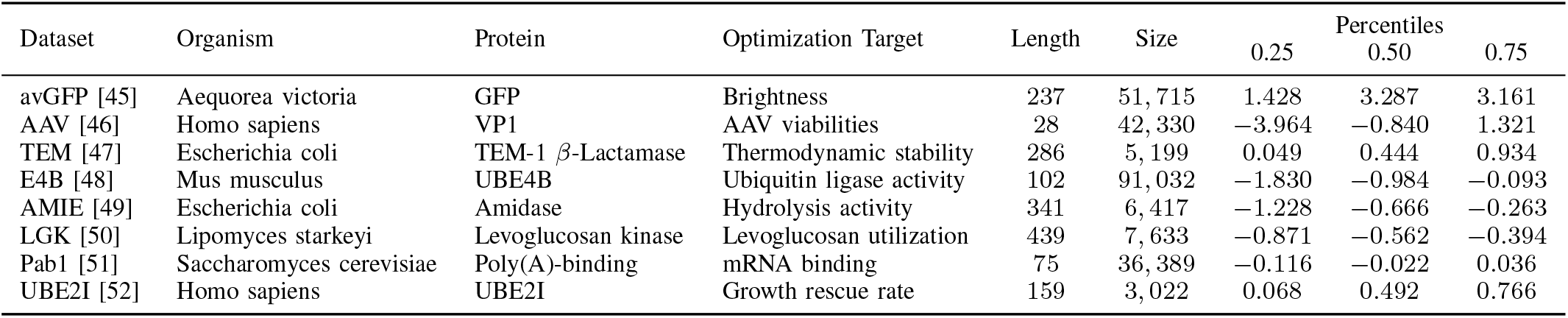
Detailed information and statistics of the eight protein datasets.

| Dataset | Organism | Protein | Optimization Target | Length | Size | 0.25 | Percentiles<br>0.50 | 0.75 |
| --- | --- | --- | --- | --- | --- | --- | --- | --- |
| avGFP [45] | Aequorea victoria | GFP | Brightness | 237 | 51,715 | 1.428 | 3.287 | 3.161 |
| AAV [46] | Homo sapiens | VP1 | AAV viabilities | 28 | 42,330 | -3.964 | -0.840 | 1.321 |
| TEM [47] | Escherichia coli | TEM-1 β-Lactamase | Thermodynamic stability | 286 | 5,199 | 0.049 | 0.444 | 0.934 |
| E4B [48] | Mus musculus | UBE4B | Ubiquitin ligase activity | 102 | 91,032 | -1.830 | -0.984 | -0.093 |
| AMIE [49] | Escherichia coli | Amidase | Hydrolysis activity | 341 | 6,417 | -1.228 | -0.666 | -0.263 |
| LGK [50] | Lipomyces starkeyi | Levoglucosan kinase | Levoglucosan utilization | 439 | 7,633 | -0.871 | -0.562 | -0.394 |
| Pab1 [51] | Saccharomyces cerevisiae | Poly(A)-binding | mRNA binding | 75 | 36,389 | -0.116 | -0.022 | 0.036 |
| UBE2I [52] | Homo sapiens | UBE2I | Growth rescue rate | 159 | 3,022 | 0.068 | 0.492 | 0.766 |

#### b) Implementation Details

The model training is conducted using a single NVIDIA A100 card, employing the Adam optimization algorithm [53] with a learning rate of 2e-4. Each dataset is randomly split into training and validation sets at a ratio of 9:1. To control the disentanglement property in the latent representation, we adopt the strategy proposed by [54] and set the expected KL values to be 20. We train the VAE model for 130 epochs and choose the best checkpoint for later inference. The experiments are run five times, and the average scores are reported. For inference, we perform gradient ascent as the warm-up phase for *T* = 500 iterations with the learning rate *α ∈* [0.001, 0.01]. The latent-based directed evolution involves *G* = 10 iterative processes, with the candidate number set to *K × B* = 128. In our implementation, we set the number of samples and beam size to *K* = 128 and *B* = 1, respectively. Finally, for the random exploration, we try multiple combinations of annealing factor *δ* = 0.1 and exploration step size *γ ∈* [1.5, 6] and report the best outcomes.

#### c) Baseline Algorithms

We compare our method against the following representative baselines: (1) **AdaLead** [55] is an advanced implementation of model-guided evolution. (2) **DyNA PPO** [43] applies proximal policy optimization to search sequences on a learned landscape model. (3) **DbAS** [56] is a probabilistic modeling framework and uses an adaptive sampling algorithm. (4) **CbAS** [17] improves on DbAS by conditioning on the desired properties. (5) **CMA-ES** [57] is a famous evolutionary search algorithm. (6) **COMs** [58] is conservative objective models for offline MBO. (7) **PEX** [27] is a model-guided sequence design algorithm using proximal exploration. (8) **GFN-AL** [59] applies GFlowNet to design biological sequences. (9) **GGS** [60] is a graph-based smoothing method to optimize protein sequences. To ensure precise evaluation, we re-execute and re-evaluate all baseline methods using the same oracle. For the implementation from (1) to (5), we employ the open-source implementation provided by [55]. Regarding other baseline methods, we utilize the codes released by their respective authors. We were unable to evaluate [28] due to unrunnable public code.

#### d) Oracles

To ensure unbiased evaluation and avoid circular use of oracles, following [61], we use two separate oracles for each fitness dataset: (1) the optimization oracle that guides the model optimization and (2) the evaluation oracle that assesses the performance of the methods. Following [27], we freeze the ESM-based encoders and fine-tune an attention 1D decoder stacked after them to predict the fitness scores. For fair comparisons, we only train the optimization oracle with the pre-trained 33-layer ESM-2 as the encoder, while using the pre-trained evaluation oracle provided by [27] to assess our method and other baselines.

#### e) Evaluation Metrics

We use three metrics defined in [62] to evaluate our method and compare with other baselines: (1) **MFS**: maximum fitness score, (2) **Diversity**, (3) **Novelty**.

The metrics are computed as follows:

- 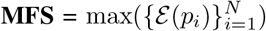,
- 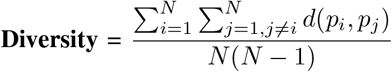,
- 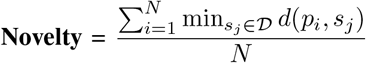,

where *d*(·,·) is the Levenshtein distance, and *𝒟* is the initial dataset (i.e., training dataset). It is crucial to emphasize that greater diversity and novelty do *not* equate to superior performance, but offer insights into the exploration and exploitation trade-offs exhibited by different methods.

### B. Results

#### a) Comparison with Baseline Algorithms

As demonstrated in Table II, our proposed LDE outperforms other algorithms in eight protein benchmarks. It is worth noting that AAV only contains sequences with up to 28 amino acids. For datasets with longer sequences exceeding 200 amino acids, using continuous latent representations demonstrably enhances LDE’s optimization capabilities. This is particularly evident on the avGFP benchmark (comprising sequences of 237 amino acids), where it achieves a statistically significant outperformance over the baselines. This suggests that LDE’s efficient exploration is particularly well-suited for tackling complex and longer protein sequences.

**TABLE II.** Maximum fitness scores across eight protein datasets. Higher values indicate better functional properties in the dataset.

|  | avGFP | AAV | TEM | E4B | AMIE | LGK | PabI | UBE2I | Average |
| --- | --- | --- | --- | --- | --- | --- | --- | --- | --- |
| $x^{wt}$ | 1.408 | -6.778 | -0.015 | 0.774 | -2.789 | -1.260 | 0.014 | -0.262 | -1.114 |
| AdaLead | 3.323 | -1.545 | 0.248 | -0.373 | -1.483 | -0.047 | 0.382 | 2.765 | 0.409 |
| DyNA PPO | 5.331 | -2.817 | 0.570 | -0.575 | -2.790 | -0.060 | 0.183 | 2.630 | 0.309 |
| DbAS | 5.141 | -2.889 | 0.493 | -0.432 | -2.093 | -0.029 | 0.228 | 2.740 | 0.395 |
| CbAS | 5.187 | -2.800 | 0.481 | -0.658 | -1.784 | -0.056 | 0.276 | 2.693 | 0.417 |
| CMA-ES | 5.125 | -3.267 | 0.590 | -0.658 | -2.790 | -0.086 | 0.254 | 2.527 | 0.212 |
| COMs | 3.544 | -3.533 | 0.472 | -0.860 | -20.182 | -0.087 | 0.156 | 2.086 | -2.300 |
| PEX | 3.796 | 2.378 | 0.252 | 4.317 | -0.364 | 0.009 | 1.326 | 3.578 | 1.911 |
| GFN-AL | 5.028 | -4.444 | 0.654 | -0.831 | -37.360 | -5.738 | 1.399 | 3.850 | -4.682 |
| GS | 3.368 | 2.442 | 1.121 | -1.147 | -3.364 | -0.972 | 0.059 | 4.101 | 0.702 |
| **LDE (ours)** | **8.058** | **2.636** | **1.745** | **5.120** | **-0.103** | **0.018** | **1.548** | **4.297** | **2.915** |

Additionally, Tables III and IV outline the diversity and novelty metrics for all methods. It is crucial to emphasize that while these results offer insights into the exploration and exploitation trade-off, they may not necessarily indicate the overall efficacy of an algorithm. The effectiveness of an algorithm depends on its specific objectives. For example, some methods aim to optimize the fitness score while maximizing diversity and novelty [28], while others may prioritize minimizing novelty [27], and some may solely focus on optimizing the fitness score alone [60]. Additionally, it’s worth mentioning that a random algorithm can achieve maximum diversity and novelty scores. This is demonstrated in the case of GFN-AL and COMs in the AMIE dataset, where they achieve high novelty and diversity scores but obtain the lowest MFS and AFS among all methods.

**TABLE III.** Diversity across eight protein datasets. This table provides insight into the exploration and exploitation trade-off among methods.

| Models | avGFP | AAV | TEM | E4B | AMIE | LGK | PabI | UBE2I | Average |
| --- | --- | --- | --- | --- | --- | --- | --- | --- | --- |
| AdaLead | 9.814 | 6.904 | 8.071 | 7.412 | 6.898 | 7.430 | 7.441 | 7.439 | 7.676 |
| DyNA PPO | 204.376 | 114.229 | 157.964 | 140.617 | 171.123 | 205.098 | 185.145 | 179.212 | 169.721 |
| DbAS | 205.717 | 115.426 | 159.524 | 142.044 | 172.539 | 206.819 | 186.761 | 180.758 | 171.199 |
| CbAS | 205.709 | 115.437 | 159.502 | 142.033 | 172.526 | 206.823 | 186.766 | 180.769 | 171.196 |
| CMA-ES | 173.365 | 96.882 | 134.633 | 119.388 | 145.151 | 174.274 | 157.098 | 152.058 | 144.106 |
| COMs | 70.319 | 45.761 | 73.191 | 68.398 | 100.731 | 126.623 | 115.242 | 109.650 | 88.739 |
| PEX | 7.048 | 4.782 | 6.692 | 6.625 | 6.388 | 7.031 | 7.012 | 7.219 | 6.600 |
| GFN-AL | 6.253 | 0.501 | 1.234 | 29.057 | 46.577 | 11.248 | 27.134 | 3.253 | 28.231 |
| GS | 4.630 | 12.712 | 7.385 | 10.577 | 14.692 | 16.151 | 16.730 | 3.011 | 10.744 |
| **LDE (ours)** | **94.646** | **1.604** | **61.652** | **4.504** | **35.762** | **108.053** | **9.040** | **14.666** | **41.806** |

**TABLE IV.** Novelty across eight datasets. this table provides insight into the exploration and exploitation trade-off among methods.

| Models | avGFP | AAV | TEM | E4B | AMIE | LGK | PabI | UBE2I | Average |
| --- | --- | --- | --- | --- | --- | --- | --- | --- | --- |
| AdaLead | 13.486 | 17.805 | 41.405 | 48.934 | 41.167 | 78.868 | 73.526 | 78.548 | 47.967 |
| DyNA PPO | 201.702 | 111.566 | 156.784 | 139.227 | 170.368 | 205.815 | 185.686 | 179.801 | 168.869 |
| DbAS | 201.838 | 111.544 | 156.858 | 139.222 | 170.066 | 205.439 | 185.366 | 179.545 | 168.735 |
| CbAS | 201.825 | 111.549 | 156.845 | 139.172 | 170.031 | 205.404 | 185.354 | 179.549 | 168.716 |
| CMA-ES | 202.155 | 111.467 | 156.968 | 139.126 | 157.701 | 193.991 | 175.414 | 170.746 | 163.446 |
| COMs | 184.177 | 98.831 | 111.931 | 101.930 | 123.590 | 150.657 | 134.298 | 129.795 | 129.401 |
| PEX | 4.323 | 1.930 | 4.461 | 4.748 | 3.031 | 8.614 | 4.356 | 4.779 | 4.530 |
| GFN-AL | 220.632 | 2.452 | 255.939 | 29.062 | 326.255 | 413.951 | 64.046 | 145.458 | 192.728 |
| GS | 3.552 | 2.061 | 4.029 | 8.081 | 9.989 | 20.990 | 8.380 | 2.862 | 7.493 |
| **LDE (ours)** | **91.863** | **0.657** | **62.508** | **2.037** | **10.423** | **65.145** | **2.875** | **126.495** | **45.250** |

#### b) Effect of Gradient Ascent and Directed Evolution

We conduct an ablation study to demonstrate how each component of our method contributes to the performance. In particular, we compare LDE with its variants, including (1) without gradient ascent (w/o GA): we do not use GA as the warm-up step for the latent-based DE and (2) without directed evolution (w/o DE): we only run gradient ascent to optimize fitness of the sequences. As shown in Table V, removing gradient ascent (i.e., w/o GA) notably reduces the performance of the evolution algorithm. Similarly, performing gradient ascent without DE only leads to sub-optimal solutions.

**TABLE V.**
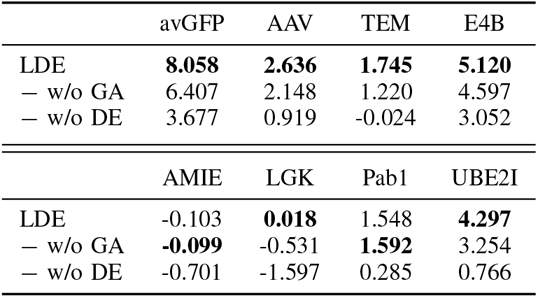
Reported maximum fitness scores of lde with/without different components on two datasets. all experiments were conducted five times using learning rate *α* = 0.002 and exploration step size *γ* = 3.

#### c) Ability to Design Low-order Mutants

Synthesizing protein sequences with minimal mutations from wild-type references offers both cost reduction [63], [64] and good biochemical performance [65], [66]. PEX [27] was built for this purpose: to discover novel proteins with low mutation count upon the wild-type sequence by incorporating distances from the wild-type sequence, along with fitness scores, into the objective function of the optimization algorithm. Since our primary objective is to design sequences with high properties and not to heavily consider mutation counts, we may not achieve mutation counts as low as those achieved by PEX in general (i.e., across the entire population). However, to showcase the ability to design low-order mutants with high fitness, we selectively choose sequences that have mutation counts close to those achieved by PEX and exhibit good fitness values. As shown in Figure 3, compared to PEX, LDE achieves lower mutation counts in AAV, avGFP, TEM, and E4B benchmarks while maintaining comparable mutation counts in AMIE and LGK. By solely focusing on maximizing protein fitness, our method produces superior performance in fitness scores across datasets. Our experiment additionally reveals that for each fitness landscape, there may be trade-offs between the number of mutations and fitness scores.

**Fig. 3.**
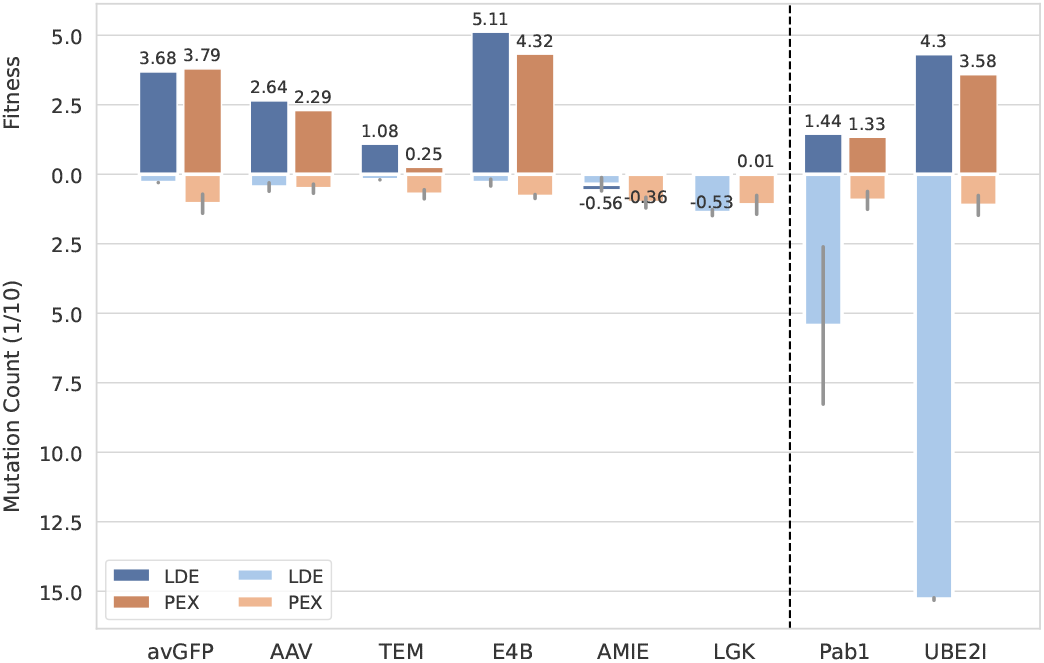
The fitness and corresponding mutation count of sequence in multiple benchmarks. For PEX, we select the best-designed sequence, while for LDE, we choose a sequence with a mutation count similar to that of PEX. The text above each bar denotes the fitness value associated with that specific bar.

#### d) Visualizations of Latent Space

Figures 4a and 4b illustrate the approximated fitness landscapes of protein sequences in the E4B family. These visualizations imply that by applying the regularization to VAEs as outlined in Equation (3), the latent representations can be organized following their fitness levels. Furthermore, as illustrated in Figure 4a, our VAEs demonstrate the capability to generate meaningful latent representations when grounded in actual fitness values for labeling these vectors. In contrast, as depicted in Figure 4b, the jointly-trained predictor *f* effectively approximates a smooth fitness landscape by predicting fitness values of latent vectors in close alignment with their true values, thereby facilitating the gradient ascent step. These observations provide a rationale for the effectiveness of our proposed latent-based directed evolution, leveraging the expressiveness power of deep learning.

**Fig. 4.**
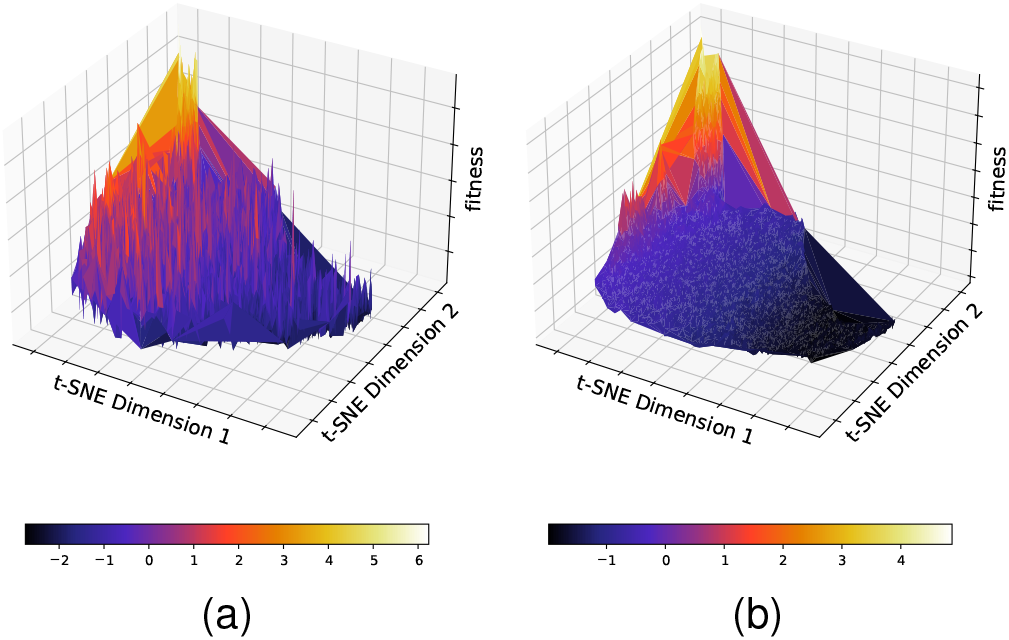
The fitness landscape approximated by regularized VAEs of E4B proteins. We compute latent representations of sequences in the validation set; then, we use t-SNE to project these latent codes into 2D spaces and visualize them with their corresponding fitness values. (a): Fitness scores are the ground truth provided by the dataset. (b): Fitness scores are predicted by the predictor *f*.

#### e) Effect of Latent Dimension

To have a comprehensive understanding of latent-based DE, we study the effect of latent dimensions on the evolution process. Table VII shows that a large latent dimension can lead to worse performance in optimizing protein fitness, while smaller latent sizes can result in sub-optimal fitness. This is an expected behavior in LSO where high dimensions may impose challenges for optimization algorithms and lower dimensions can not encode sufficient information about the input data. Furthermore, we observe that novelty and diversity are independent of good fitness and vary specifically for different fitness landscapes.

#### f) Autoregressive vs. Non-autoregressive

In addition to the autoregressive decoder utilized in LDE, we report the performance of a non-autoregressive decoder implemented following the architecture proposed by [34]. This decoder comprises four 1-dimensional convolutional layers with ReLU activations and batch normalization layers are incorporated between convolutional layers, except for the final layer. As outlined in Table VI, it is observed that the autoregressive decoder consistently outperforms the non-autoregressive decoder across all tasks. This paves the way for further study on how different types of decoders affect the latent-based evolutionary algorithms.

**TABLE VI.** Maximum fitness scores on eight protein datasets of two decoder versions of lde.

|  | avGFP | AAV | TEM | E4B | AMIE | LGK | PabI | UBE2I | Average |
| --- | --- | --- | --- | --- | --- | --- | --- | --- | --- |
| Non-autoregressive LDE | 3.733 | 1.368 | 1.095 | 3.123 | -0.558 | -0.005 | 0.089 | 2.976 | 1.430 |
| (Autoregressive) LDE | **8.058** | **2.636** | **1.745** | **5.120** | **-0.103** | **0.018** | **1.548** | **4.297** | **3.204** |

**TABLE VII.** Reported metrics of lde with different latent sizes on two datasets. all experiments were conducted five times using learning rate *α* = 0.002 and exploration step size *γ* = 3.

| Dataset | Latent Dim | Fitness | Novelty | Diversity |
| --- | --- | --- | --- | --- |
| E4B<br>(L = 102) | 128 | 4.433 | 2.040 | 5.322 |
|  | 256 | 4.597 | 0.615 | 0.968 |
|  | 320 | **5.120** | 2.038 | 4.504 |
|  | 512 | 3.052 | 0.862 | 0.968 |
| TEM<br>(L = 286) | 128 | 1.408 | 17.109 | 44.906 |
|  | 256 | 1.220 | 267.013 | 1.783 |
|  | 320 | **1.745** | 62.508 | 61.652 |
|  | 512 | -0.024 | 269.992 | 0.044 |

## VI. Conclusion

We present Latent-based Directed Evolution (LDE), a novel method that combines directed evolution with gradient ascent in a regularized VAE latent space to efficiently optimize and design protein sequences. This approach leverages deep representation learning capabilities of generative models to significantly speed up the evolutionary process, achieving superior results compared to traditional methods solely operating in sequence space. LDE holds significant promise for accelerating protein engineering and drug discovery efforts, and we invite further research on integrating it with in vitro protein characterization for real-world validation.

## References

[1] S. Biswas, G. Khimulya, E. C. Alley, K. M. Esvelt, and G. M. Church, “Low-n protein engineering with data-efficient deep learning,” Nature Methods, vol. 18, no. 4, p. 389–396, Apr. 2021. [Online]. Available: 10.1038/s41592-021-01100-y

[2] R. J. Fox, S. C. Davis, E. C. Mundorff, L. M. Newman, V. Gavrilovic, S. K. Ma, L. M. Chung, C. Ching, S. Tam, S. Muley, J. Grate, J. Gruber, J. C. Whitman, R. A. Sheldon, and G. W. Huisman, “Improving catalytic function by prosar-driven enzyme evolution,” Nature Biotechnology, vol. 25, no. 3, p. 338–344, Feb. 2007. [Online]. Available: 10.1038/nbt1286

[3] H. D. Lagassé, A. Alexaki, V. L. Simhadri, N. H. Katagiri, W. Jankowski, Z. E. Sauna, and C. Kimchi-Sarfaty, “Recent advances in (therapeutic protein) drug development,” F1000Research, vol. 6, p. 113, Feb. 2017. [Online]. Available: 10.12688/f1000research.9970.1

[4] N. Go, “Theoretical studies of protein folding,” Annual Review of Biophysics and Bioengineering, vol. 12, no. 1, p. 183–210, Jun. 1983. [Online]. Available: 10.1146/annurev.bb.12.060183.001151

[5] C. Chothia, “Principles that determine the structure of proteins,” Annual Review of Biochemistry, vol. 53, no. 1, p. 537–572, Jun. 1984. [Online]. Available: 10.1146/annurev.bi.53.070184.002541

[6] S. Wright, “The roles of mutation, inbreeding, crossbreeding and selection in evolution,” Proceedings of the XI International Congress of Genetics, vol. 8, pp. 209–222, 1932.

[7] J. Maynard Smith, “Natural selection and the concept of a protein space,” Nature, vol. 225, no. 5232, p. 563–564, Feb. 1970. [Online]. Available: 10.1038/225563a0

[8] F. H. Arnold, “Directed evolution: Creating biocatalysts for the future,” Chemical Engineering Science, vol. 51, no. 23, pp. 5091–5102, 1996. [Online]. Available: https://www.sciencedirect.com/science/article/pii/S0009250996002886

[9] F. H. Arnold, “Design by directed evolution,” Accounts of chemical research, vol. 31, no. 3, pp. 125–131, 1998.

[10] F. H. Arnold and A. A. Volkov, “Directed evolution of biocatalysts,” Current opinion in chemical biology, vol. 3, no. 1, pp. 54–59, 1999.

[11] Y. Qiu, J. Hu, and G.-W. Wei, “Cluster learning-assisted directed evolution,” Nature Computational Science, vol. 1, no. 12, p. 809–818, Dec. 2021. [Online]. Available: 10.1038/s43588-021-00168-y

[12] B. J. Wittmann, Y. Yue, and F. H. Arnold, “Informed training set design enables efficient machine learning-assisted directed protein evolution,” Cell Systems, vol. 12, no. 11, pp. 1026–1045.e7, Nov. 2021. [Online]. Available: 10.1016/j.cels.2021.07.008

[13] Y. Qiu and G.-W. Wei, “Clade 2.0: Evolution-driven cluster learning-assisted directed evolution,” Journal of Chemical Information and Modeling, vol. 62, no. 19, p. 4629–4641, Sep. 2022. [Online]. Available: 10.1021/acs.jcim.2c01046

[14] P. Emami, A. Perreault, J. Law, D. Biagioni, and P. St. John, “Plug and play directed evolution of proteins with gradient-based discrete mcmc,” Machine Learning: Science and Technology, vol. 4, no. 2, p. 025014, Apr. 2023. [Online]. Available: 10.1088/2632-2153/accacd

[15] D. H. Brookes, A. Aghazadeh, and J. Listgarten, “On the sparsity of fitness functions and implications for learning,” Proceedings of the National Academy of Sciences, vol. 119, no. 1, p. e2109649118, 2022.

[16] C. Dallago, J. Mou, J. Mou, K. Johnston, B. Wittmann, N. Bhattacharya, S. Goldman, A. Madani, and K. Yang, “Flip: Benchmark tasks in fitness landscape inference for proteins,” in Proceedings of the Neural Information Processing Systems Track on Datasets and Benchmarks, J. Vanschoren and S. Yeung, Eds., vol. 1. Curran, 2021. [Online]. Available: https://datasets-benchmarks-proceedings.neurips.cc/paperfiles/paper/2021/file/2b44928ae11fb9384c4cf38708677c48-Paper-round2.pdf

[17] D. Brookes, H. Park, and J. Listgarten, “Conditioning by adaptive sampling for robust design,” in Proceedings of the 36th International Conference on Machine Learning, ser. Proceedings of Machine Learning Research, K. Chaudhuri and R. Salakhutdinov, Eds., vol. 97. PMLR, 09–15 Jun 2019, pp. 773–782. [Online]. Available: https://proceedings.mlr.press/v97/brookes19a.html

[18] A. Kumar and S. Levine, “Model inversion networks for modelbased optimization,” in Advances in Neural Information Processing Systems, H. Larochelle, M. Ranzato, R. Hadsell, M. Balcan, and H. Lin, Eds., vol. 33. Curran Associates, Inc., 2020, pp. 5126–5137. [Online]. Available: https://proceedings.neurips.cc/paperfiles/paper/2020/file/373e4c5d8edfa8b74fd4b6791d0cf6dc-Paper.pdf

[19] Z. Lin, H. Akin, R. Rao, B. Hie, Z. Zhu, W. Lu, N. Smetanin, R. Verkuil, O. Kabeli, Y. Shmueli, A. dos Santos Costa, M. Fazel-Zarandi, T. Sercu, S. Candido, and A. Rives, “Evolutionary-scale prediction of atomic-level protein structure with a language model,” Science, vol. 379, no. 6637, pp. 1123–1130, 2023. [Online]. Available: https://www.science.org/doi/abs/10.1126/science.ade2574

[20] T. A. Hopf, J. B. Ingraham, F. J. Poelwijk, C. P. I. Schärfe, M. Springer, C. Sander, and D. S. Marks, “Mutation effects predicted from sequence co-variation,” Nature Biotechnology, vol. 35, no. 2, p. 128–135, Jan. 2017. [Online]. Available: 10.1038/nbt.3769

[21] A. J. Riesselman, J. B. Ingraham, and D. S. Marks, “Deep generative models of genetic variation capture the effects of mutations,” Nature Methods, vol. 15, no. 10, p. 816–822, Sep. 2018. [Online]. Available: 10.1038/s41592-018-0138-4

[22] Y. Luo, G. Jiang, T. Yu, Y. Liu, L. Vo, H. Ding, Y. Su, W. W. Qian, H. Zhao, and J. Peng, “Ecnet is an evolutionary context-integrated deep learning framework for protein engineering,” Nature Communications, vol. 12, no. 1, Sep. 2021. [Online]. Available: 10.1038/s41467-021-25976-8

[23] R. Rao, N. Bhattacharya, N. Thomas, Y. Duan, P. Chen, J. Canny, P. Abbeel, and Y. Song, “Evaluating protein transfer learning with tape,” in Advances in Neural Information Processing Systems, H. Wallach, H. Larochelle, A. Beygelzimer, F. d’Alché-Buc, E. Fox, and R. Garnett, Eds., vol. 32. Curran Associates, Inc., 2019. [Online]. Available: https://proceedings.neurips.cc/paperfiles/paper/2019/file/37f65c068b7723cd7809ee2d31d7861c-Paper.pdf

[24] E. C. Alley, G. Khimulya, S. Biswas, M. AlQuraishi, and G. M. Church, “Unified rational protein engineering with sequence-based deep representation learning,” Nature Methods, vol. 16, no. 12, p. 1315–1322, Oct. 2019. [Online]. Available: 10.1038/s41592-019-0598-1

[25] J. Meier, R. Rao, R. Verkuil, J. Liu, T. Sercu, and A. Rives, “Language models enable zero-shot prediction of the effects of mutations on protein function,” in Advances in Neural Information Processing Systems, M. Ranzato, A. Beygelzimer, Y. Dauphin, P. Liang, and J. W. Vaughan, Eds., vol. 34. Curran Associates, Inc., 2021, pp. 29 287–29 303. [Online]. Available: <https://proceedings.neurips.cc/paperfiles/paper/2021/file/f51338d736f95dd42427296047067694-Paper.pdf

[26] C. Hsu, H. Nisonoff, C. Fannjiang, and J. Listgarten, “Learning protein fitness models from evolutionary and assay-labeled data,” Nature Biotechnology, vol. 40, no. 7, p. 1114–1122, Jan. 2022. [Online]. Available: 10.1038/s41587-021-01146-5

[27] Z. Ren, J. Li, F. Ding, Y. Zhou, J. Ma, and J. Peng, “Proximal exploration for model-guided protein sequence design,” in Proceedings of the 39th International Conference on Machine Learning, ser. Proceedings of Machine Learning Research, K. Chaudhuri, S. Jegelka, L. Song, C. Szepesvari, G. Niu, and S. Sabato, Eds., vol. 162. PMLR, 17–23 Jul 2022, pp. 18 520–18 536. [Online]. Available: https://proceedings.mlr.press/v162/ren22a.html

[28] Z. Song and L. Li, “Importance weighted expectation-maximization for protein sequence design,” in Proceedings of the 40th International Conference on Machine Learning, ser. Proceedings of Machine Learning Research, A. Krause, E. Brunskill, K. Cho, B. Engelhardt, S. Sabato, and J. Scarlett, Eds., vol. 202. PMLR, 23–29 Jul 2023, pp. 32 349–32 364. [Online]. Available: https://proceedings.mlr.press/v202/song23g.html

[29] E. T. Farinas, T. Bulter, and F. H. Arnold, “Directed enzyme evolution,” Current opinion in biotechnology, vol. 12, no. 6, pp. 545–551, 2001.

[30] C. A. Tracewell and F. H. Arnold, “Directed enzyme evolution: climbing fitness peaks one amino acid at a time,” Current opinion in chemical biology, vol. 13, no. 1, pp. 3–9, 2009.

[31] F. H. Arnold, “Directed evolution: Bringing new chemistry to life,” Angewandte Chemie International Edition, vol. 57, no. 16, pp. 4143–4148, 2018. [Online]. Available: https://onlinelibrary.wiley.com/doi/abs/10.1002/anie.201708408

[32] T. T. Tran and T. S. Hy, “Protein design by directed evolution guided by large language models,” bioRxiv, 2023. [Online]. Available: https://www.biorxiv.org/content/early/2023/11/29/2023.11.28.568945

[33] R. Gómez-Bombarelli, J. N. Wei, D. Duvenaud, J. M. Hernández-Lobato, B. Sánchez-Lengeling, D. Sheberla, J. Aguilera-Iparraguirre, T. D. Hirzel, R. P. Adams, and A. Aspuru-Guzik, “Automatic chemical design using a data-driven continuous representation of molecules,” ACS Central Science, vol. 4, no. 2, pp. 268–276, 2018, pMID: 29532027. [Online]. Available: 10.1021/acscentsci.7b00572

[34] E. Castro, A. Godavarthi, J. Rubinfien, K. Givechian, D. Bhaskar, and S. Krishnaswamy, “Transformer-based protein generation with regularized latent space optimization,” Nature Machine Intelligence, vol. 4, pp. 1–12, 09 2022.

[35] M. Lee, L. F. Vecchietti, H. Jung, H. Ro, M. Cha, and H. M. Kim, “Protein sequence design in a latent space via model-based reinforcement learning,” 2023. [Online]. Available: https://openreview.net/forum?id=OhjGzRE5N6o

[36] S. Stanton, W. Maddox, N. Gruver, P. Maffettone, E. Delaney,P. Greenside, and A. G. Wilson, “Accelerating Bayesian optimization for biological sequence design with denoising autoencoders,” in Proceedings of the 39th International Conference on Machine Learning, ser. Proceedings of Machine Learning Research, K. Chaudhuri, S. Jegelka, L. Song, C. Szepesvari, G. Niu, and S. Sabato, Eds., vol. 162. PMLR, 17–23 Jul 2022, pp. 20 459–20 478. [Online]. Available: https://proceedings.mlr.press/v162/stanton22a.html

[37] T. Chen, P. Vure, R. Pulugurta, and P. Chatterjee, “AMP-diffusion: Integrating latent diffusion with protein language models for antimicrobial peptide generation,” in NeurIPS 2023 Generative AI and Biology (GenBio) Workshop, 2023. [Online]. Available: https://openreview.net/forum?id=145TM9VQhx

[38] K. E. Johnston, C. Fannjiang, B. J. Wittmann, B. L. Hie, K. K. Yang, and Z. Wu, “Machine learning for protein engineering,” arXiv preprint 2305.16634, 2023.

[39] Y. Wang, P. Xue, M. Cao, T. Yu, S. T. Lane, and H. Zhao, “Directed evolution: Methodologies and applications,” Chemical Reviews, vol. 121, no. 20, pp. 12 384–12 444, Oct 2021. [Online]. Available: 10.1021/acs.chemrev.1c00260

[40] O. Sener and S. Savarese, “Active learning for convolutional neural networks: A core-set approach,” in International Conference on Learning Representations, 2018. [Online]. Available: https://openreview.net/forum?id=H1aIuk-RW

[41] W. Jin, R. Barzilay, and T. Jaakkola, “Junction tree variational autoencoder for molecular graph generation,” in Proceedings of the 35th International Conference on Machine Learning, ser. Proceedings of Machine Learning Research, J. Dy and A. Krause, Eds., vol. 80. PMLR, 10–15 Jul 2018, pp. 2323–2332. [Online]. Available: https://proceedings.mlr.press/v80/jin18a.html

[42] D. P. Kingma, S. Mohamed, D. Jimenez Rezende, and M. Welling, “Semi-supervised learning with deep generative models,” in Advances in Neural Information Processing Systems, Z. Ghahramani, M. Welling, C. Cortes, N. Lawrence, and K. Weinberger, Eds., vol. 27. Curran Associates, Inc., 2014. [Online]. Available: https://proceedings.neurips.cc/paperfiles/paper/2014/file/d523773c6b194f37b938d340d5d02232-Paper.pdf

[43] C. Angermueller, D. Dohan, D. Belanger, R. Deshpande, K. Murphy, and L. Colwell, “Model-based reinforcement learning for biological sequence design,” in International Conference on Learning Representations, 2020. [Online]. Available: https://openreview.net/forum?id=HklxbgBKvr

[44] X. Chen, D. P. Kingma, T. Salimans, Y. Duan, P. Dhariwal, J. Schulman Sutskever, and P. Abbeel, “Variational lossy autoencoder,” in International Conference on Learning Representations, 2017. [Online]. Available: https://openreview.net/forum?id=BysvGP5ee

[45] K. S. Sarkisyan, D. A. Bolotin, M. V. Meer, D. R. Usmanova, A. S. Mishin, G. V. Sharonov, D. N. Ivankov, N. G. Bozhanova, M. S. Baranov, O. Soylemez, N. S. Bogatyreva, P. K. Vlasov, E. S. Egorov, M. D. Logacheva, A. S. Kondrashov, D. M. Chudakov, E. V. Putintseva, I. Z. Mamedov, D. S. Tawfik, K. A. Lukyanov, and F. A. Kondrashov, “Local fitness landscape of the green fluorescent protein,” Nature, vol. 533, no. 7603, pp. 397–401, May 2016. [Online]. Available: 10.1038/nature17995

[46] D. H. Bryant, A. Bashir, S. Sinai, N. K. Jain, P. J. Ogden, P. F. Riley, G. M. Church, L. J. Colwell, and E. D. Kelsic, “Deep diversification of an aav capsid protein by machine learning,” Nature Biotechnology, vol. 39, no. 6, pp. 691–696, Jun 2021. [Online]. Available: 10.1038/s41587-020-00793-4

[47] E. Firnberg, J. W. Labonte, J. J. Gray, and M. Ostermeier, “A Comprehensive, High-Resolution Map of a Gene’s Fitness Landscape,” Molecular Biology and Evolution, vol. 31, no. 6, pp. 1581–1592, 02 2014. [Online]. Available: 10.1093/molbev/msu081

[48] L. M. Starita, J. N. Pruneda, R. S. Lo, D. M. Fowler, H. J. Kim, J. B. Hiatt, J. Shendure, P. S. Brzovic, S. Fields, and R. E. Klevit, “Activity-enhancing mutations in an e3 ubiquitin ligase identified by high-throughput mutagenesis,” Proceedings of the National Academy of Sciences, vol. 110, no. 14, pp. E1263–E1272, 2013. [Online]. Available: https://www.pnas.org/doi/abs/10.1073/pnas.1303309110

[49] E. E. Wrenbeck, L. R. Azouz, and T. A. Whitehead, “Single-mutation fitness landscapes for an enzyme on multiple substrates reveal specificity is globally encoded,” Nature Communications, vol. 8, no. 1, p. 15695, Jun 2017. [Online]. Available: 10.1038/ncomms15695

[50] J. R. Klesmith, J.-P. Bacik, R. Michalczyk, and T. A. Whitehead, “Comprehensive sequence-flux mapping of a levoglucosan utilization pathway in e. coli,” ACS Synthetic Biology, vol. 4, no. 11, p. 1235–1243, Sep. 2015. [Online]. Available: 10.1021/acssynbio.5b00131

[51] D. Melamed, D. L. Young, C. E. Gamble, C. R. Miller, and S. Fields, “Deep mutational scanning of an rrm domain of the saccharomyces cerevisiae poly(a)-binding protein,” RNA, vol. 19, no. 11, p. 1537–1551, Sep. 2013. [Online]. Available: 10.1261/rna.040709.113

[52] J. Weile, S. Sun, A. G. Cote, J. Knapp, M. Verby, J. C. Mellor, Y. Wu, C. Pons, C. Wong, N. van Lieshout, F. Yang, M. Tasan, G. Tan, S. Yang, D. M. Fowler, R. Nussbaum, J. D. Bloom, M. Vidal, D. E. Hill, P. Aloy, and F. P. Roth, “A framework for exhaustively mapping functional missense variants,” Molecular Systems Biology, vol. 13, no. 12, p. 957, 2017. [Online]. Available: https://www.embopress.org/doi/abs/10.15252/msb.20177908

[53] D. P. Kingma and J. Ba, “Adam: A method for stochastic optimization,” in 3rd International Conference on Learning Representations, ICLR 2015, San Diego, CA, USA, May 7-9, 2015, Conference Track Proceedings, Y. Bengio and Y. LeCun, Eds., 2015. [Online]. Available: http://arxiv.org/abs/1412.6980

[54] H. Shao, S. Yao, D. Sun, A. Zhang, S. Liu, D. Liu, J. Wang, and T. Abdelzaher, “ControlVAE: Controllable variational autoencoder,” in Proceedings of the 37th International Conference on Machine Learning, ser. Proceedings of Machine Learning Research, H. D. III and A. Singh, Eds., vol. 119. PMLR, 13–18 Jul 2020, pp. 8655–8664. [Online]. Available: https://proceedings.mlr.press/v119/shao20b.html

[55] S. Sinai, R. Wang, A. Whatley, S. Slocum, E. Locane, and E. D. Kelsic, “Adalead: A simple and robust adaptive greedy search algorithm for sequence design,” CoRR, vol. abs/2010.02141, 2020. [Online]. Available: https://arxiv.org/abs/2010.02141

[56] D. H. Brookes and J. Listgarten, “Design by adaptive sampling,” 2020.

[57] N. Hansen and A. Ostermeier, “Completely derandomized self-adaptation in evolution strategies,” Evolutionary Computation, vol. 9, no. 2, pp. 159–195, 2001.

[58] B. Trabucco, A. Kumar, X. Geng, and S. Levine, “Conservative objective models for effective offline model-based optimization,” in Proceedings of the 38th International Conference on Machine Learning, ser. Proceedings of Machine Learning Research, M. Meila and T. Zhang, Eds., vol. 139. PMLR, 18–24 Jul 2021, pp. 10 358–10 368. [Online]. Available: https://proceedings.mlr.press/v139/trabucco21a.html

[59] M. Jain, E. Bengio, A. Hernandez-Garcia, J. Rector-Brooks, B. F. Dossou, C. A. Ekbote, J. Fu, T. Zhang, M. Kilgour, D. Zhang et al., “Biological sequence design with gflownets,” in International Conference on Machine Learning. PMLR, 2022, pp. 9786–9801.

[60] A. Kirjner, J. Yim, R. Samusevich, S. Bracha, T. S. Jaakkola, R. Barzilay, and I. R. Fiete, “Improving protein optimization with smoothed fitness landscapes,” in The Twelfth International Conference on Learning Representations, 2023.

[61] S. Kolli, A. X. Lu, X. Geng, A. Kumar, and S. Levine, “Data-driven optimization for protein design: Workflows, algorithms and metrics,” in ICLR2022 Machine Learning for Drug Discovery, 2022. [Online]. Available: https://openreview.net/forum?id=Dc5J-bcEGW5

[62] M. Jain, E. Bengio, A. Hernandez-Garcia, J. Rector-Brooks, B. F. P. Dossou, C. A. Ekbote, J. Fu, T. Zhang, M. Kilgour, D. Zhang, L. Simine, P. Das, and Y. Bengio, “Biological sequence design with GFlowNets,” in Proceedings of the 39th International Conference on Machine Learning, ser. Proceedings of Machine Learning Research, K. Chaudhuri, S. Jegelka, L. Song, C. Szepesvari, G. Niu, and S. Sabato, Eds., vol. 162. PMLR, 17–23 Jul 2022, pp. 9786–9801. [Online]. Available: https://proceedings.mlr.press/v162/jain22a.html

[63] H. H. Hogrefe, J. Cline, G. L. Youngblood, and R. M. Allen, “Creating randomized amino acid libraries with the quikchange® multi site-directed mutagenesis kit,” Biotechniques, vol. 33, no. 5, pp. 1158–1165, 2002.

[64] P. E. Carrigan, P. Ballar, and S. Tuzmen, “Site-directed mutagenesis,” Disease Gene Identification: Methods and Protocols, pp. 107–124, 2011.

[65] J. R. Klesmith, J.-P. Bacik, E. E. Wrenbeck, R. Michalczyk, and T. A. Whitehead, “Trade-offs between enzyme fitness and solubility illuminated by deep mutational scanning,” Proceedings of the National Academy of Sciences, vol. 114, no. 9, pp. 2265–2270, 2017.

[66] C. L. Araya and D. M. Fowler, “Deep mutational scanning: assessing protein function on a massive scale,” Trends in biotechnology, vol. 29, no. 9, pp. 435–442, 2011.

