## Appendix for "Latent-based Directed Evolution accelerated by Gradient Ascent for Protein Sequence Design"

### I. Variational Autoencoders

Variational autoencoders (VAEs) [1] is a class of generative model wherein each data sample $x$ is generated by a generative distribution $p_\theta(x|z)$, parameterized by $\theta$, that conditions on the unseen low dimensional latent variable $z \in \mathbb{R}^d$, which can be sampled from a prior distribution $p(z)$ (usually assumed as Gaussian). In other words, $\theta$ is trained to maximize the marginal log-likelihood $\log p_\theta(x) := \log \int_z p_\theta(x|z)p(z)dz$. However, this is intractable as all the configurations of the latent variable $z$ must be examined during the optimization. To address this issue, [1] proposed to use amortized variational inference to approximate the true posterior distribution $p_\theta(z|x)$ via a variational distribution $q_\phi(z|x)$ with learnable parameters $\phi$. They, instead, optimize the evidence of the lower bound (ELBO) of $\log p(x)$, which is written as:

$$\log p(x) \geq \mathbb{E}_{q_\phi(z|x)}(\log p_\theta(x|z)) - \beta D_{\text{KL}}(q_\phi(z|x)||p(z)). \quad (1)$$

From the perspective of autoencoding, $\phi$ and $\theta$ are regarded as an encoder (inference network) and decoder (generative network), respectively. The encoder $\phi$ maps the high-dimensional input variable $x$ to the low-dimensional latent variable $z$ within the latent space. In particular, the mean $\mu_\phi(x)$ and log-variance $\log \sigma_\phi(x)$ of the posterior $q_\phi(z|x)$ are computed, and $z$ is sampled from $q_\phi(z|x) \sim \mathcal{N}(\mu_\phi(x), \sigma_\phi(x)^2)$. Finally, the decoder $\theta$ maps $z$ back to the input space.

Popular VAE applications often involve a trade-off between reconstruction accuracy and some other application-specific goals (e.g., to produce *new and original* candidates), effectively manipulated through KL-divergence. ControlVAE [2] combines control theory [3] with the basic VAE to stabilize the KL-divergence to a desired value. It designs a PI controller to dynamically tune the weight $\beta$ in the Equation (1) by using the actual KL-divergence as feedback during training as follows:

$$\beta(t) = K_p \sigma(-e(t)) - K_i \sum_{j=0}^{t} e(j) + \beta_{\min}, \quad (2)$$

where $e(t) = C - D_{\text{KL}}(q_\phi(z|x)||p(z))(t)$, which is the difference between desired KL-divergence $C$ and the actual one at training step $t$; $\sigma(\cdot)$ is a sigmoid function; $\beta_{\min}$ is a constant; $K_p$ and $K_i$ are positive co-efficients for the P term and I term respectively.

† Corresponding author: Truong Son Hy.
* Nhat Khang Ngo and Thanh V. T. Tran contribute equally to this work.
Nhat Khang Ngo, Thanh V. T. Tran, and Viet Thanh Duy Nguyen are with FPT Software AI Center, Hanoi, Vietnam
Truong Son Hy is with Indiana State University, Terre Haute, USA

### II. Data Descriptions

*a) Green Fluorescent Proteins (**avGFP**):* Derived from Aequorea victoria, Green Fluorescent proteins (GFPs) are capable of manifesting vivid green fluorescence upon exposure to light within the blue to ultraviolet spectrum. These proteins are commonly employed as biosensors for detecting gene expressions and protein locations. Our objective is to design sequences with enhanced capabilities as gene delivery vectors, quantified by AAV viabilities. The magnitude of the search space encompasses $20^{28}$ possibilities.

*b) Adeno-associated Viruses (**AAV**):* The engineer of a 28-amino acid segment (position 561–588) within the VP1 protein, situated in the capsid of the Adeno-associated virus, has garnered significant interest in the realm of machine learning-guided design. We aim to design more capable sequences as gene delivery vectors measured by AAV viabilities. The size of the search space is $20^{28}$.

*c) Aliphatic Amide Hydrolase (**AMIE**):* The enzyme encoded by amiE, known as Amidase, holds industrial relevance and is d erived from Pseudomonas aeruginosa. Our objective is to optimize amidase sequences that result in enhanced enzyme activities, defining a search space comprising $20^{341}$ sequences.

*d) Ubiquitination Factor Ube4b (**E4B**):* The ubiquitination factor Ube4b plays a pivotal role in cellular waste degradation through interactions with ubiquitin and other proteins. Our emphasis is on the enhancement of E4B enzyme activity within the specified landscape, which encompasses a search space of $20^{102}$.

*e) Levoglucosan Kinase (**LGK**):* Levoglucosan kinase catalyzes the conversion of Levoglucosan (LG) to the glycolytic intermediate glucose-6-phosphate through an ATP-dependent reaction. The target is to optimize LGK protein sequences with improved enzyme activity. The size of search space is $20^{439}$.

*f) Poly(A)-binding Protein (**Pab1**):* Pab1 utilizes the RNA recognition motif (RRM) to bind to multiple adenosine monophosphates (poly-A). The aim is to design sequences with higher binding fitness to multiple adenosine monophos-

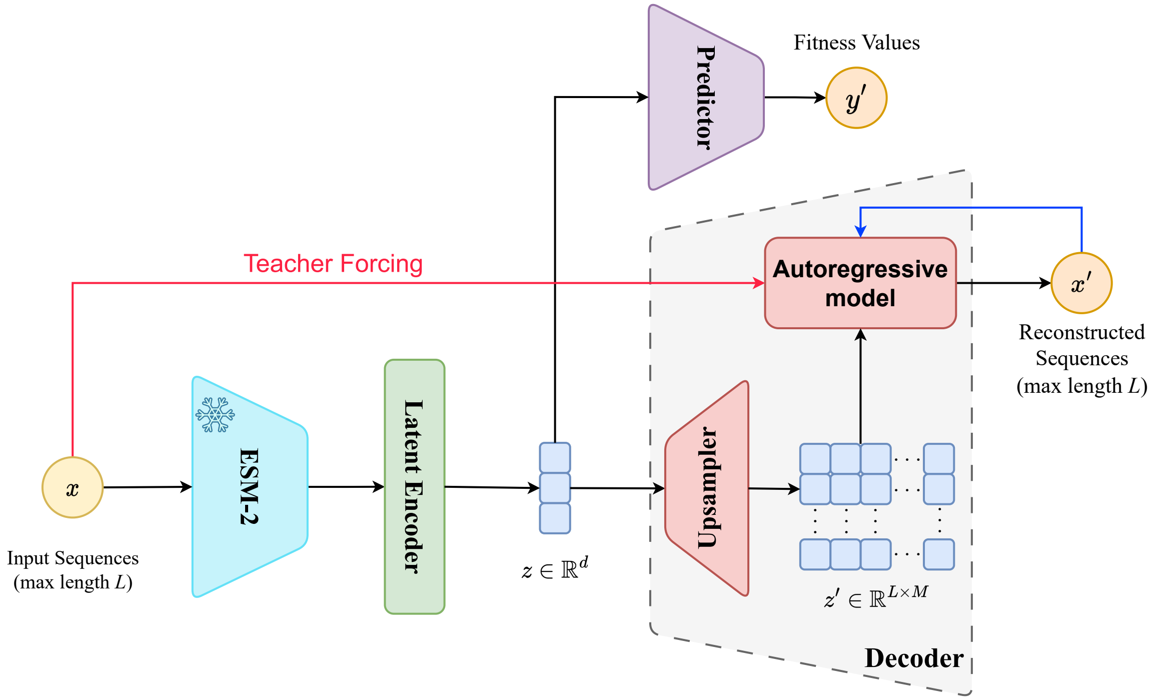


Fig. 1: Schematic illustration of the VAE model used in the study.

phates. The search space size is $20^{75}$ on a segment of the wild-type sequence.

*g) TEM-1 $\beta$-Lactamase (**TEM**):* The investigation of the TEM-1 $\beta$-Lactamase protein's resistance to penicillin antibiotics in E. coli is a subject of extensive scrutiny, aiming to comprehend mutational impacts and the associated fitness landscape. The optimization objective involves suggesting sequences with elevated thermodynamic stability compared to the wild-type TEM-1 within a search space of magnitude $20^{286}$.

*h) SUMO E2 conjugase (**UBE2I**):* The utilization of variants for the functional mapping of human genomes holds considerable significance in both scientific research and clinical treatment. The search space encompasses a size of $20^{159}$ within a segment of the wild-type sequence.

### III. Implementation Details

#### A. VAEs' Architecture

In this section, we go into detail regarding the architecture of the VAE used in our study. As mentioned in the main text, our regularized VAE consists of an encoder, a predictor, and a decoder.

*a) Encoder:* Figure 1 depicts that the encoder incorporates a pre-trained ESM-2 [4] followed by a latent encoder to compute the latent representation $z$. In our study, we leverage the powerful representation of the pre-trained 30-layer ESM-2 by making it the encoder of our model. Given an input sequence $x = \langle x_0, x_1, \cdots, x_L \rangle$, where $x_i \in \mathcal{V}$, the transformer-based ESM-2 computes representations for each token $x_i$ in $x$, resulting in a token-level hidden representation $H = \langle h_0, h_1, \cdots, h_L \rangle, h_i \in \mathbb{R}^{d_h}$. We calculate the global representation $h \in \mathbb{R}^{d_h}$ of $x$ via a weighted sum of its tokens:

$$h = \sum_{i=1}^{L} \frac{\omega^T \exp(h_i)}{\sum_{i=1}^{L} \omega^T \exp(h_i)} h_i. \quad (3)$$

here, $\omega$ is a learnable global attention vector. Then, two multi-layer perceptrons (MLPs) are used to compute $\mu = \text{MLP}_1(h)$ and $\log \sigma = \text{MLP}_2(h)$, where the latent dimension is $d$. Finally, a latent representation $z \in \mathbb{R}^d$ is sampled from $\mathcal{N}(\mu, \sigma^2)$, which is further proceeded to the decoder to reconstruct the sequence $\hat{x}$. We use an auxiliary MLP as a **fitness predictor** that maps the latent $z$ to the fitness score, i.e. $y' = \text{MLP}(z)$. The hidden dimensional size of the predictor is set to 512, with the dropout of 0.2. We set $d_h = 1280$ and $d = 320$ for the main experiments in our study.

*b) Decoder:* Inspired from [5], we construct our decoder as the combination of two components: an 'upsampler' component comprising 3 layers of transposed convolutions with stride of 2 to upsample the latent vector $z$ to a sequence matching the output sequence's length; and an autoregressive component consisting of an attention-based GRU [6], [7] with 512 units. To cope with the optimization difficulties reported when training VAE with powerful autoregressive decoders [8], [5], we follow [8] by applying $40\%$ dropout to the amino acid context supplied as input to the GRU during training. This encourages the network to depend on the information conveyed through the upsampled latent code along with the conditional information in the masked amino acid sequence to make predictions. Additionally, we apply teacher forcing [9] with a ratio of $50\%$ for faster convergence.

**Algorithm 1** Active Learning with Latent-Based DE

**Input:** a VAE $M = (\phi, \theta, f)$, training dataset $\mathcal{D}_t$, number of rounds $N$, optimization oracle $\mathcal{O}$, number of epochs $n$.

Train $M$ on $\mathcal{D}_t$
$\mathcal{D}_0 \leftarrow \emptyset$
**for** $i = 1$ **to** $N$ **do**
  Run Algorithm 1 (main text) with oracle $\mathcal{O}$ and $M$ to find population $\mathcal{P} = \{(x, \mathcal{O}(y)) | x \in \mathcal{V}^L, y \in \mathbb{R}\}$.
  $\mathcal{D}_i \leftarrow \mathcal{D}_{i-1} \cup$ RemoveDuplicate($\mathcal{P}$)
  Update $M$ on the data $\mathcal{D}_i$ in $n$ epochs
**end for**

**Return** $P_G$

TABLE I: Comparison of optimization and evaluation oracles on the max fitness scores across eight protein benchmarks.

| | avGFP | AAV | TEM | E4B | AMIE | LGK | Pab1 | UBE2I |
|---|---|---|---|---|---|---|---|---|
| Opt. Oracle $\mathcal{O}$ | 15.266 | 2.736 | 5.986 | 5.594 | 0.327 | 0.939 | 0.786 | 7.405 |
| Eval. Oracle $\mathcal{E}$ | 8.058 | 2.636 | 1.745 | 5.120 | -0.103 | 0.018 | 1.548 | 4.297 |

#### B. Oracles

We establish the optimization oracle $\mathcal{O}(\cdot)$, utilized for latent-based directed evolution, by leveraging features generated by the pre-trained 33-layer ESM-2 [4] with a dimension of 1280. Subsequently, we fine-tune an Attention1D model to predict fitness values based on these representations. As for the evaluation oracle $\mathcal{E}(\cdot)$, which acts as the "ground-truth" evaluator, we employ the trained oracle provided by [10]. This evaluation oracle combines the pre-trained ESM-1b [11] with a Attention1D model with a dimension of 512, and it is used to assess all methods.

#### C. Training Configurations

As mentioned in the Section I, we employ the ControlVAE mechanism to prevent KL vanishing and enhance the diversity of generated data during training. Across all tasks, we configure the coefficients $K_p$ and $K_i$ of the P term and I term to 0.01 and 0.0001, respectively. For the LGK benchmark, the desired KL-divergence $C$ is set to 40, while for all other tasks, $C = 20$ is utilized. The batch size for each task is determined to be as large as possible, as long as the total steps in one epoch for each task are higher than 100.

### IV. Additional Results

#### A. Differences between optimization oracle and evaluation oracle

In Table I, we present a comparison of results for designs using the optimization oracle $\mathcal{O}(\cdot)$ (Opt. Oracle) and the evaluation oracle $\mathcal{E}(\cdot)$ (Eval. Oracle). The table demonstrates that, with the exception of Pab1, all other benchmarks experienced a decrease in performance when using the evaluation oracle. This decline is expected, as different oracle architectures result in different approximate fitness scores. Therefore, as we optimize the results based on the optimization oracle and solely utilize the evaluation oracle to assess the final performance of the method, the score produced by the optimization oracle should be higher.

#### B. Active Learning

Protein fitness datasets are often fragmented due to the cost of wet lab experiments [12], and even then, they only capture a small protein of real-world protein behavior. This limitation can trap machine learning models, which learn from training samples, in local optima, hindering their generalizability and accuracy [13]. To overcome this issue, in this work, we propose using active learning to update the fitness landscape approximated by our regularized VAEs. Indeed, Algorithm 1 demonstrates our method in detail.

In particular, we perform an outer active learning loop with $N$ rounds to iteratively update the latent space, as well as the simulated landscape produced by the encoder $\phi$ and the latent fitness predictor $f$ as mentioned in the main text. For each round $i$, we fine-tune the VAE model $M$ with the dataset $\mathcal{D}_{i-1}$. The fine-tuned model is then used in Algorithm I in the main text to explore sequences with higher fitness scores. In our study, we avoid the circular use of oracles by using the optimization oracle $\mathcal{O}(\cdot)$ during the optimization process to guide the exploration, and the evaluation oracle $\mathcal{E}(\cdot)$ is used to evaluate of the final population. We remove all duplicated samples in $\mathcal{P}$ and use them as training samples $\mathcal{D}_i$ for the next round in our algorithm.

To validate our proposed method, we conduct additional experiments on the four smallest benchmark datasets, each containing fewer than $8,000$ training data points: TEM, AMIE, LGK, and UBE2I. For this demonstration, we set the number of active learning rounds $N$ to 10 and decrease the number of directed evolution iterations $G$ to 5. Additionally, in each active learning loop, the regularized VAE $M$ is fine-tuned in 30 epochs. As outlined in Table II, the non-autoregressive LDE demonstrates improved performance when combined with active learning. These results empirically validate our hypothesis and confirm the efficacy of our proposed method.

### V. Experiments with Another Oracle

To sufficiently demonstrate the efficacy of our method, we have trained a new evaluation oracle to assess the perfor-

TABLE II: Max fitness scores on four smallest protein datasets.

| Model | TEM | AMIE | LGK | UBE2I |
|---|---|---|---|---|
| LDE | 1.095 | -0.558 | -0.005 | 2.976 |
| − w/ active learning | **2.167** | **-0.015** | **0.022** | **3.698** |

mance of both baseline methods and our proposed approach. Specifically, we have substituted the ESM-1b backbone of the evaluation oracle $\mathcal{E}$ with the 150M-parameter ESM-2 model and increased the hidden size of Attention1D module from 512 to 640. The datasets used are the same for both oracles. The results of most recent methods are reported in Table III

### References


[1] D. P. Kingma, S. Mohamed, D. Jimenez Rezende, and M. Welling, "Semi-supervised learning with deep generative models," *Advances in neural information processing systems*, vol. 27, 2014.

[2] H. Shao, S. Yao, D. Sun, A. Zhang, S. Liu, D. Liu, J. Wang, and T. Abdelzaher, "Controlvae: Controllable variational autoencoder," in *International conference on machine learning*. PMLR, 2020, pp. 8655–8664.

[3] K. J. Åström and T. Hägglund, *Advanced PID control*. ISA-The Instrumentation, Systems and Automation Society, 2006.

[4] Z. Lin, H. Akin, R. Rao, B. Hie, Z. Zhu, W. Lu, N. Smetanin, R. Verkuil, O. Kabeli, Y. Shmueli *et al.*, "Evolutionary-scale prediction of atomic-level protein structure with a language model," *Science*, vol. 379, no. 6637, pp. 1123–1130, 2023.

[5] S. Semeniuta, A. Severyn, and E. Barth, "A hybrid convolutional variational autoencoder for text generation," in *Proceedings of the 2017 Conference on Empirical Methods in Natural Language Processing*, M. Palmer, R. Hwa, and S. Riedel, Eds. Copenhagen, Denmark: Association for Computational Linguistics, Sep. 2017, pp. 627–637. [Online]. Available: https://aclanthology.org/D17-1066

[6] K. Cho, B. van Merriënboer, C. Gulcehre, D. Bahdanau, F. Bougares, H. Schwenk, and Y. Bengio, "Learning phrase representations using RNN encoder–decoder for statistical machine translation," in *Proceedings of the 2014 Conference on Empirical Methods in Natural Language Processing (EMNLP)*, A. Moschitti, B. Pang, and W. Daelemans, Eds. Doha, Qatar: Association for Computational Linguistics, Oct. 2014, pp. 1724–1734. [Online]. Available: https://aclanthology.org/D14-1179

[7] T. Luong, H. Pham, and C. D. Manning, "Effective approaches to attention-based neural machine translation," in *Proceedings of the 2015 Conference on Empirical Methods in Natural Language Processing*, L. Màrquez, C. Callison-Burch, and J. Su, Eds. Lisbon, Portugal: Association for Computational Linguistics, Sep. 2015, pp. 1412–1421. [Online]. Available: https://aclanthology.org/D15-1166

[8] S. R. Bowman, L. Vilnis, O. Vinyals, A. Dai, R. Jozefowicz, and S. Bengio, "Generating sentences from a continuous space," in *Proceedings of the 20th SIGNLL Conference on Computational Natural Language Learning*, S. Riezler and Y. Goldberg, Eds. Berlin, Germany: Association for Computational Linguistics, Aug. 2016, pp. 10–21. [Online]. Available: https://aclanthology.org/K16-1002

[9] R. J. Williams and D. Zipser, "A learning algorithm for continually running fully recurrent neural networks," *Neural computation*, vol. 1, no. 2, pp. 270–280, 1989.

[10] Z. Ren, J. Li, F. Ding, Y. Zhou, J. Ma, and J. Peng, "Proximal exploration for model-guided protein sequence design," in *International Conference on Machine Learning*. PMLR, 2022, pp. 18 520–18 536.

[11] A. Rives, J. Meier, T. Sercu, S. Goyal, Z. Lin, J. Liu, D. Guo, M. Ott, C. L. Zitnick, J. Ma *et al.*, "Biological structure and function emerge from scaling unsupervised learning to 250 million protein sequences," *Proceedings of the National Academy of Sciences*, vol. 118, no. 15, p. e2016239118, 2021.

[12] C. Dallago, J. Mou, K. E. Johnston, B. Wittmann, N. Bhattacharya, S. Goldman, A. Madani, and K. K. Yang, "FLIP: Benchmark tasks in fitness landscape inference for proteins," in *Thirty-fifth Conference on Neural Information Processing Systems Datasets and Benchmarks Track (Round 2)*, 2021. [Online]. Available: https://openreview.net/forum?id=p2dMLEwL8tF

[13] D. Brookes, H. Park, and J. Listgarten, "Conditioning by adaptive sampling for robust design," in *International conference on machine learning*. PMLR, 2019, pp. 773–782.

TABLE III: Max fitness scores on different oracle across most recent baselines.

| Method | avGFP | AAV | TEM | E4B | AMIE | LGK | Pab1 | UBE2I | Average |
|---|---|---|---|---|---|---|---|---|---|
| GFN-AL | 6.21 | -2.30 | 2.46 | 1.40 | -0.14 | 0.03 | **1.80** | 1.78 | 1.41 |
| PEX | 3.32 | 2.78 | 0.62 | 3.85 | **0.01** | 0.76 | 0.69 | 1.49 | 1.69 |
| GGS | 3.79 | **3.77** | 1.24 | 0.04 | -1.06 | 0.24 | 0.47 | 3.42 | 1.49 |
| LDE | **7.88** | 3.17 | **3.05** | **6.43** | -0.01 | **0.89** | 0.66 | **11.96** | **4.25** |
